# Trend and pattern of antimicrobial resistance in molluscan *Vibrio* species sourced to Canadian estuaries

**DOI:** 10.1101/305821

**Authors:** Swapan K. Banerjee, Jeffrey M. Farber

**Affiliations:** Bureau of Microbial Hazards, Food Directorate, Health Canada 251 promenade Sir Frederick Banting Driveway, Ottawa, ON, Canada K1A 0K9; Department of Food Science, Canadian Research Institute for Food Safety (CRIFS) University of Guelph, Food Science Bldg, Guelph, ON, Canada N1G 2W1

**Keywords:** *Vibrio* spp., antimicrobial resistance, surveillance, molluscan shellfish

## Abstract

Emergence of antimicrobial resistance (AMR) in foodborne bacteria is a growing concern worldwide. AMR surveillance is a key element in understanding the implications resulting from the use of antibiotics for therapeutic as well as prophylactic needs. The emergence and spread of AMR in foodborne human pathogens is an indirect health hazard. This surveillance study reports the trend and pattern of AMR detected in *Vibrio* species isolated from molluscs harvested in Canada, between 2006 and 2012, against 19 commonly used antibiotics. Five common antibiotics, ampicillin, cephalothin, erythromycin, kanamycin and streptomycin, predominantly contributed to AMR including multi-drug resistance (MDR) in the molluscan *Vibrio* spp. isolated in 2006. A prospective follow-up analysis of these drugs showed a declining trend in the frequency of MDR/AMR-*Vibrio* spp. in subsequent years until 2012. The observed decline appears to have been influenced by the specific downturn in resistance to the aminoglycosides, kanamycin and streptomycin. Frequently observed MDR/AMR*-Vibrio* spp. in seafood is a potential health concern associated with seafood consumption. Our surveillance study provided an indication of the antibiotics that challenged the marine bacteria, sourced to Canadian estuaries, during and/or prior to the study period.

## INTRODUCTION

Halophilic ‘non-cholera’ *Vibrio* species, particularly *V. parahaemolyticus* and *V. alginolyticus*, are natural inhabitants of the estuarine waters around the world, making them suitable for monitoring ecosystem challenges, such as the impact of using antibiotics and chemicals in aquaculture, farms and clinical facilities. Use of antibiotics in aquaculture and animal husbandry for prophylaxis and growth promotion has the potential to affect human and animal health. Of various ecological environments, the estuarine ecosystem provides an interactive opportunity for the marine bacteria to encounter washouts containing residues of antibiotics from farmlands and hospitals, making estuaries the most dynamic natural habitat to complement or confirm events of the nearby lands (1). Antibiotic resistance genes predate our use of antibiotics and the first direct evidence was reported after analysis of DNA sequences recovered from Late Pleistocene permafrost sediments to indicate that antibiotic resistance is an ancient, naturally occurring phenomenon detectable in the environment (2). Antimicrobial selective pressure is particularly high in hospitals where inpatients, for example, in 20-30% of cases in Europe, receive a dose of antibiotics during hospitalization and treatment (3). In Canada, over 150 billion litres of untreated or under-treated sewage are dumped into the waterways every year (4), and the amount of polluting materials, including antibiotics, is not always controlled. Some of the marine bacteria, including halophilic *Vibrio* spp., survive antibiotic exposure by acquiring antimicrobial resistance (AMR) mechanisms. The exchange of resistance determinants between the aquatic and the terrestrial environments can result from the transfer and interaction of AMR bacteria between the two environments. The consumption of fishery products is on the rise among health conscious consumers and seafood-lovers, which may be partly the reason for the increase in seafood-borne illnesses.

Antibiotics have been a critical public health tool since the discovery of penicillin in 1928 by Alexander Fleming. The introduction of the drug as a therapeutic agent in the 1940s saved the lives of millions of people around the world (5). Thereafter, antimicrobial drugs of various other classes, subclasses and their subsequent generations were introduced to the therapeutic regimen to fight bacterial infections. However, within a few years of introduction of the antibiotics, the emergence of drug resistance in bacteria capable of inactivating drugs, or blocking their action or entry into the cell, evolved to reverse the gains of the beneficial discoveries. Hence, many important drug choices for the treatment of bacterial infections have become limited and can yield unpredictable results. At the same time, not much progress has been made in the discovery of new antibiotics and challenges remain for the future (6). The Centers for Disease Control and Prevention (CDC) reported that an estimated two million illnesses and 23,000 deaths are caused by drug-resistant bacteria in the USA alone (7). The European Surveillance of Antimicrobial Consumption (ESAC) reported that macrolide use was the main driver behind the emergence of macrolide resistance in streptococci, and documented a positive correlation between the level of antibiotic resistance and the level of antibiotic consumption (8). Effective intervention with antibiotics and the proper stewardship of antibiotic use, will require evidence-based knowledge regarding the status of AMR at regional and national levels. The World Health Organisation (WHO) proposed and initiated a Global Action Plan on AMR in 2015 (9); one of the strategic objectives of the Plan was to strengthen the evidence base through enhanced surveillance and analyses. Thereafter, the Global AMR Surveillance System (GLASS) was adopted to enable standardised and comparable data to be collected, analysed and shared between countries for advocacy and preventative action. Of the many gaps identified by the WHO reports (9, 10), information on *Vibrio* spp. of marine origin with AMR profile and their impact on public health is lacking. Prospective follow-up of bacterial AMR patterns, as observed in the molluscan *Vibrio* spp., studied over a long period of time, can provide information on potential trends occurring from the widespread use of antibiotics and its direct or indirect impacts on human health. Similar AMR data on selected bacteria from terrestrial animals are reported by the Canadian Integrated Program for Antimicrobial Resistance Surveillance (CIPARS) to understand trends in antimicrobial use and resistance, for the benefit of intervention and therapeutic strategies (11, 12). Evidence of resistance to clinically important antimicrobials among selected bacteria isolated from retail meats in Canada’ has shown a wide variation over time and across regions covering farmlands and poultry (13). Clinically significant AMR *Vibrio* spp. can be ingested by humans through seafood, and can potentially lead to serious consequences. This study reports our findings to determine the extent and dynamics of AMR detected predominantly among six halophilic *Vibrio* species (*V. alginolyticus, V. parahaemolyticus, V. cholerae, V. vulnificus, V. fluvialis* and *Vibrio* spp.) used as indicator bacteria isolated from molluscs, such as oysters, mussels or clams harvested in Canadian estuaries from 2006 to 2012.

## RESULTS

### Resistance and virulence profile were detected in surveillance

Halophilic *Vibrio* species, which are accumulated by filter-feeding molluscs asymptomatically in the estuarine habitat, displayed resistance to some of the 19 antibiotics used in this study including multidrug-resistance (MDR, three or more antibiotics). In addition, potential virulence factor genes *(tdh*, and/or *trh)* were detected by PCR in some of the (AMR) isolates. Overall 343 mollusc samples were tested during the summer months of each year from 2006 to 2012, yielding six different *Vibrio* species with various frequencies of detection (Table 1). Multiple isolates and/or species belonging to the genus *Vibrio* were captured from most of the samples. In total, 1021 strains of *Vibrio* species (Table 2) were isolated during the study period yielding an average of three isolates per sample. Only 4.9% of the isolates tested sensitive to all 19 drugs. A total of 319 (31.2%) and 588 (57.6%) of the isolates were identified as *V. parahaemolyticus* and *V. alginolyticus*, respectively, of which 47 *V. parahaemolyticus* (14.7%) tested positive for either, or rarely both, of the virulence markers *(tdh* and *trh)* by PCR, and 3 *V. alginolyticus* (0.5%) tested positive for *trh* marker by PCR. These potential pathogens harboured variable AMR profiles, including some (n=17) *V. parahaemolyticus* isolates showing MDR. Out of the 17 MDR isolates, four were *tdh^+^ trh^+^* and the remainder were *tdh trh^+^* by PCR.

**Table 1.** Total number of molluscan samples tested during May to October of each year from Canadian harvest sites on either coast. The data from Atlantic and Pacific sites are shown separately, including the species identified from each region and the number of samples testing positive for each. Six types of *Vibrio* species such as, *V. parahaemolyticus* (Vp), *V. alginolyticus* (Va), *V. vulnificus* (Vv), *V. cholerae* (Vc), *V. fluvialis* (Vf), and unidentified *Vibrio* species (Vspp) were detected. In 2006, Va, Vf and Vspp were not isolated (shown as N/I), therefore, the total number of samples was adjusted to calculate the average frequency (% positive) for those species.

| YEAR | Total<br>No. of<br>samples<br>tested | East coast (Atlantic) molluscs |  |  |  |  |  |  | West coast (Pacific) molluscs |  |  |  |  |  |  |
| --- | --- | --- | --- | --- | --- | --- | --- | --- | --- | --- | --- | --- | --- | --- | --- |
|  |  | No. of<br>samples | No. of samples positive (%) for |  |  |  |  |  | No. of<br>samples | No. of samples positive (%) for |  |  |  |  |  |
|  |  |  | Vp | Va | Vv | Vc | Vf | Vspp |  | Vp | Va | Vv | Vc | Vf | Vspp |
| 2006 | 31 | 15 | 8 | N/I | 0 | 0 | N/I | N/I | 16 | 10 | N/I | 0 | 0 | N/I | N/I |
| 2007 | 37 | 18 | 8 | 13 | 2 | 1 | 0 | 0 | 19 | 12 | 19 | 0 | 0 | 0 | 0 |
| 2008 | 46 | 22 | 8 | 19 | 8 | 0 | 3 | 4 | 24 | 15 | 22 | 4 | 0 | 4 | 6 |
| 2009 | 59 | 27 | 10 | 26 | 6 | 0 | 3 | 5 | 32 | 15 | 32 | 6 | 1 | 7 | 6 |
| 2010 | 62 | 29 | 16 | 29 | 7 | 1 | 1 | 6 | 33 | 18 | 32 | 1 | 1 | 4 | 7 |
| 2011 | 54 | 23 | 12 | 22 | 5 | 0 | 3 | 3 | 31 | 12 | 27 | 3 | 1 | 2 | 10 |
| 2012 | 54 | 23 | 21 | 23 | 5 | 0 | 2 | 6 | 31 | 23 | 31 | 4 | 2 | 2 | 7 |
| **Sum =** | **343** | **157** | **83**<br>**(53%)** | **132**<br>**(93%)** | **33**<br>**(21%)** | **2**<br>**(1%)** | **12**<br>**(8%)** | **24**<br>**(17%)** | **186** | **105**<br>**(56%)** | **163**<br>**(96%)** | **18**<br>**(10%)** | **5**<br>**(3%)** | **19**<br>**(11%)** | **36**<br>**(21%)** |

**Table 2.**
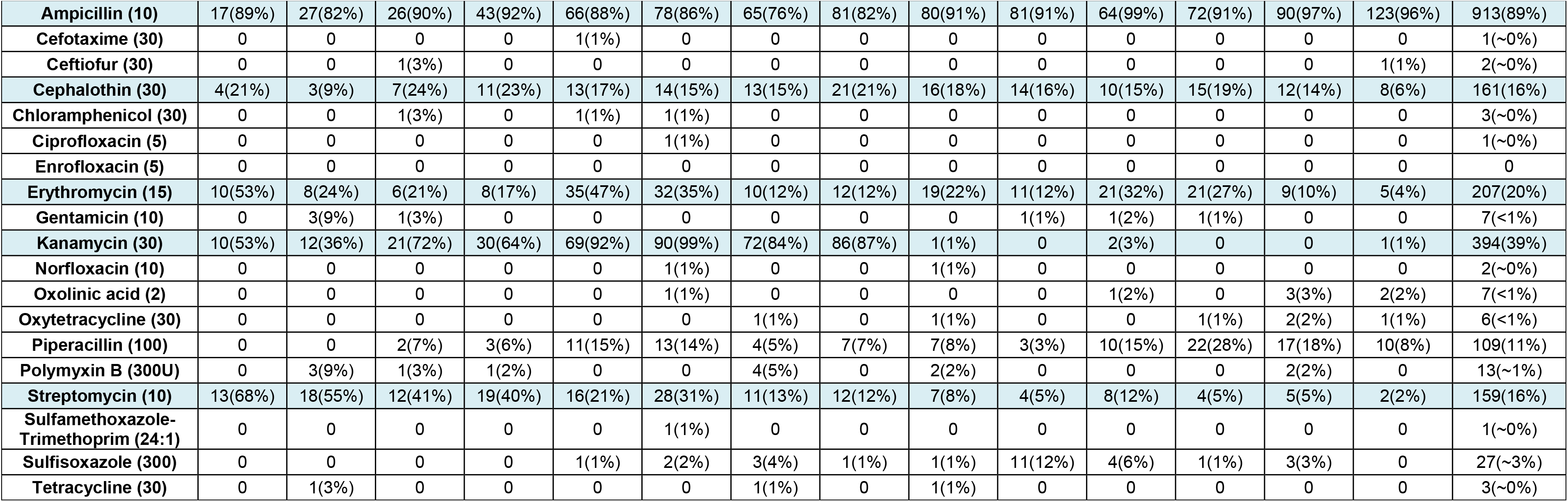
*Comparison of antimicrobial resistance (AMR) and its frequency (% in brackets) detected in marine Vibrio spp. isolated from harvest sites in the Atlantic (E) and Pacific (W)* coasts of Canada during 2006 to 2012 are shown. The five antibiotics contributing the most to AMR patterns and trends are highlighted. The number of confirmed isolates (n) by region and year are shown.

In addition to the major species, 114 strains belonging to other *Vibrio* spp. were identified (11% of total isolates) as follows: 28 *V. vulnificus*, 24 *V. fluvialis*, 7 *V. cholerae* and 55 unidentified *Vibrio* spp. showing good identification at the genus level. All of these isolates, including the 7 choleragenic vibrios of marine origin were tested for AMR and included in the analysis (Figs. 1 and 2). It should be noted here that the two major species (*V. parahaemolyticus* and *V. alginolyticus*) displayed similar AMR patterns by site, and by year (data not shown). Many isolates (n=294; 29%) of *Vibrio* species were resistant to multiple antibiotics (3 or more = MDR), but a declining trend in MDR was observed between 2008 and 2012 (Fig. 3). Resistance to 6 antibiotics was detected in three *V. parahaemolyticus* isolates, one (S191-10, British Columbia [BC], Pacific coast) in 2007 and two (S222-7, BC, Pacific coast, and S240-8, Quebec, Atlantic coast) in 2008, showing resistance to ampicillin, piperacillin, cephalothin, erythromycin, kanamycin and streptomycin. Similarly, two *V. alginolyticus* isolates (S219-1, Nova Scotia [NS], Atlantic coast, and S230-5, BC, Pacific coast) showed resistance to the same six drugs in 2008, and another isolate (S280-5, NS, Atlantic coast) in 2009 displayed a different profile in which resistance to sulfisoxazole, tetracycline and oxytetracycline was detected, while there was no resistance to piperacillin, erythromycin and streptomycin. Resistance to ampicillin and kanamycin were common to all six isolates mentioned above. The MDR frequency of the halophilic *Vibrio* spp. declined from 60% in 2008 to 7% in 2012 (Fig. 3).

**Fig 1.**
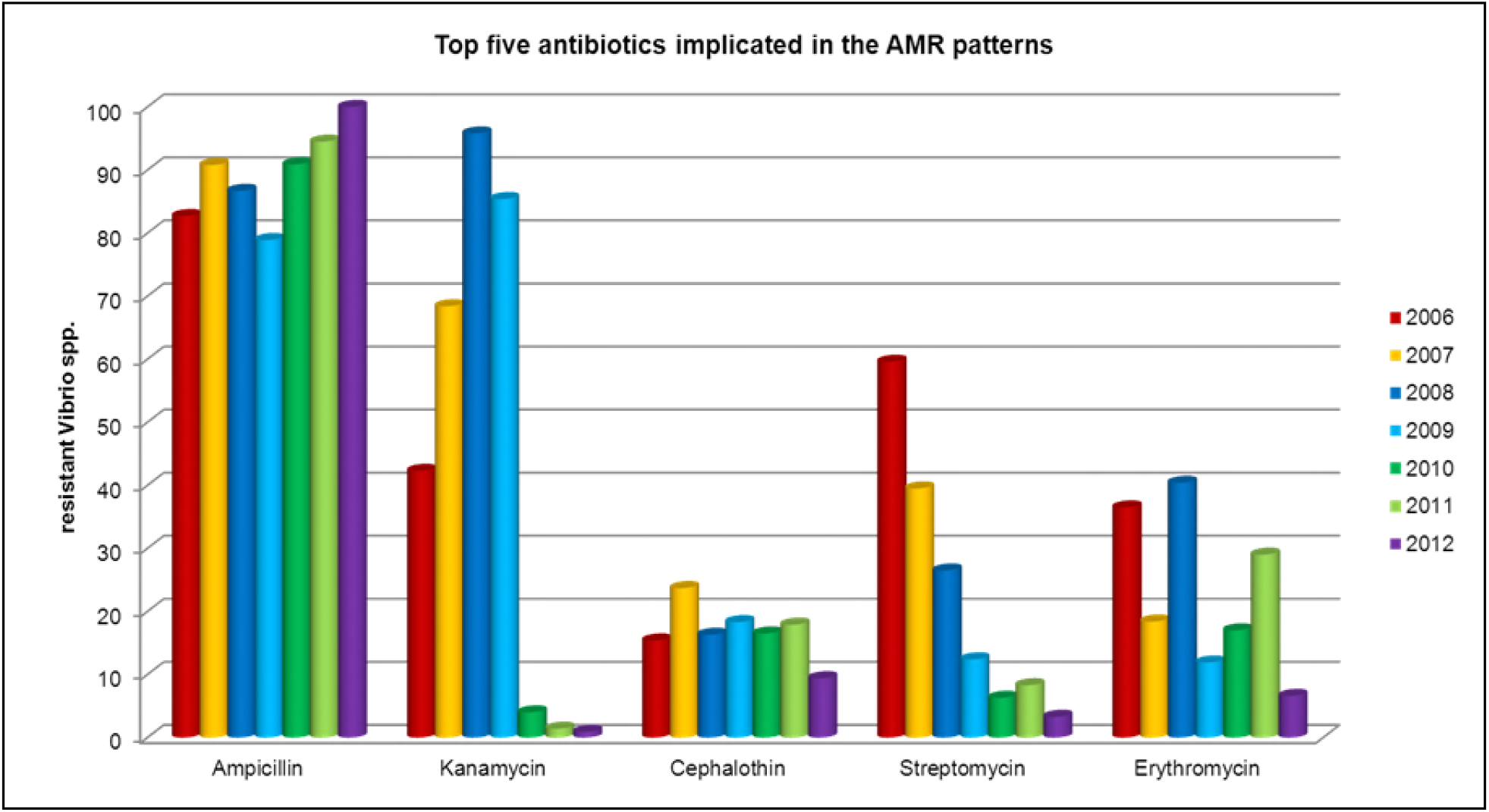
The top five antibiotics to which resistance was detected among the halophilic *Vibrio* spp. isolated seasonally from Canadian molluscan shellfish, between 2006 and 2012. The reduction in frequency of resistance over time was significant for the aminoglycosides, kanamycin and streptomycin.

**Fig 2.**
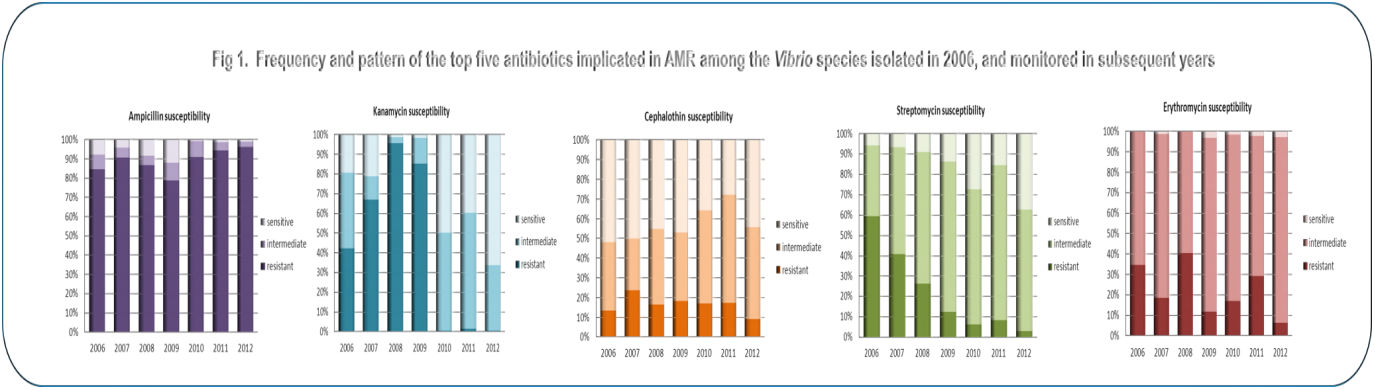
Detection frequency and complete susceptibility responses of halophilic *Vibrio* spp. to the five leading antibiotics between 2006 and 2012. While resistance to kanamycin and streptomycin declined over time, susceptibility to the other drugs increased, as shown by the darkest (resistant), lighter (intermediate) and lightest (susceptible) colors in the bars.

**Fig 3.**
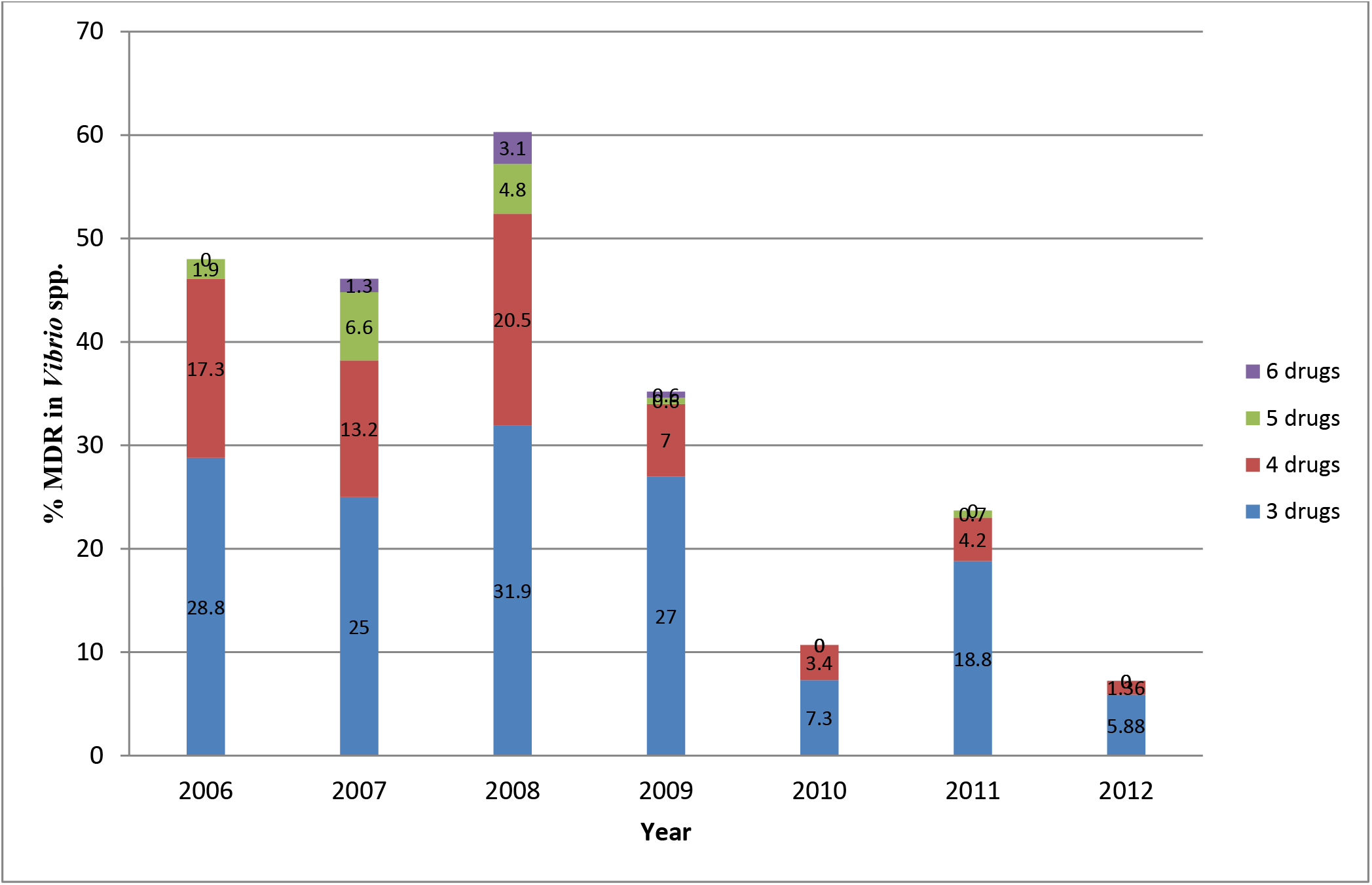
Multi-drug resistance (MDR, 3 antibiotics and more) detected among halophilic *Vibrio* spp. isolated from molluscan shellfish harvested in Canada from 2006 to 2012.

### Resistance trend observed

The top five antibiotics contributing to resistance profiles in 2006 were monitored during the period from 2006 to 2012, and a significant drop in resistance to the aminoglycosides, kanamycin and streptomycin, occurred among the *Vibrio* spp. (Figs.1 and 2). Resistance to ampicillin was detected at a very high frequency, ranging from 76% to 99% with an average of 89% during the study period (Table 1), and the difference between East and West coast data on ampicillin detected yearly was not significantly different by Student’s *t*-test (paired, twotailed, p=0.47), and the variance F-test (two-tailed, p=0.44). Similar tests with other important antibiotics also showed no significant differences between East and West coast data, e.g., in the case of kanamycin and streptomycin, the p-values were 0.42 and 0.37 (t-test), and 0.93 and 0.82 (F-test), respectively. Resistance to kanamycin showed an upward trend from 2006 to 2009, sharply declined thereafter and remained low in frequency until the end of the study period in 2012 (Figs.1 and 2). The frequency of resistance to streptomycin showed a decreasing trend during this period (Table 1). Amongst other clinically significant antibiotics, fluoroquinolones rarely contributed to AMR (Table 1).

## DISCUSSION

Since the predominant species (*V. alginolyticus* and *V. parahaemolyticus*) of *Vibrio* isolated from wild or cultured molluscs did not show any significant differences in their AMR patterns over the study period, it was presumed that the halophilic vibrios, including the lesser prevalent species, like *V. vulnificus, V. fluvialis, V. cholerae* and unidentified *Vibrio* spp., isolated from the same source during the study period, would respond more or less similarly to the exposed challenges from contaminating antibiotics. Although proto-genomic profiles of the species may vary to some extent, for our purpose, the genus *Vibrio* was used collectively as the indicator bacteria for determining the AMR pattern captured by the molluscan *Vibrio* species from the coastal environments of Canada.

The observed trend of a drop in AMR/MDR in molluscan *Vibrio* species between 2006 and 2012, could be an indication of the variability in the presence and levels of antibiotics in the Canadian estuaries. This trend could have resulted from unplanned stewardship practised for aminoglycosides during the study period, due to a preference for other antibiotics or due to the easy availability and publicized effectiveness of new drugs on the market. Available data on defined daily doses (DDD) per 1000 inhabitant-days for parenteral antibiotics purchased by hospitals in Canada from 2001 to 2011 was reported on earlier (14), and the average DDD values for aminoglycosides was 0.05 in 2006, and dropped to 0.03 in 2011 (40% drop). Similar values for first generation cephalosporins, including cephalothin, were between 0.11 and 0.12 [14]. In the USA, 42 antibiotics belonging to 18 drug classes were approved for use in food-producing animals and actively marketed in 2009 (15). Kanamycin was absent from the list of five aminoglycosides (15), and our data showing a sudden drop in resistance to kanamycin from 2010 to 2012, coincides with the drop in drug usage in the USA which shares coastal waters with Canada. A higher frequency of resistance to streptomycin (85%) and kanamycin (57%) was also observed in halophilic vibrios isolated from seafood in Italy in 2001 (16), probably indicating that these two antibiotics were also used in Italy and surrounding areas at that time. Another contemporary European (ESAC) surveillance study (8) reported that antibiotic use, defined by daily doses/1000 inhabitants/day was high in southern European countries (Greece, France and Italy), which also tends to support the Italian report (16). Therefore, emergence of AMR/MDR seems to be directly proportional to drug usage in or around the region.

The combined presence of virulence markers and AMR in some *Vibrio* strains implies a potential emerging hazard for consumers of raw or under-cooked seafood, particularly molluscan shellfish. It is difficult to establish whether those *Vibrio* strains that are both MDR and carry virulence genes, may be more hazardous for humans. Detailed analysis of strains and patients involved in *Vibrio* outbreaks will be useful in the future to generate a larger database to understand this complex issue.

A multi-pronged strategy consisting of infection control practices, surveillance and antibiotic stewardship has been recommended to slow the increasing rate of AMR [17]. D’Costa et al. (18) reported that the level and diversity of the environmental resistome is likely to be higher and much more extensive than detected in their surveillance study using soil microorganisms. Therefore, it is possible that innovation in the discovery of novel antibiotics may not be a solution to the AMR issue, as, with time, AMR could also develop in bacteria exposed to these novel antibiotics. To combat AMR, a process can be used whereby one alternates the usage of an antibiotic, or a class of antibiotics, in a cyclic manner. As a result, the targeted antibiotics can be made effective for periods of time separated by phases of complete prohibition, which would need to be determined more precisely following more objective studies and supportive data from other temperate regions of the world. Thus, a strategy of cyclic regulation of medically-important drugs could possibly be used for antibiotic stewardship programs at the international level.

This study has presented for the first time, a detailed analysis of the AMR profiles of molluscan *Vibrio* spp. isolated from molluscs grown in Canadian estuaries. As a result, the data presented here will be a useful addition to the repository of data on AMR resistance in Canada from surveillance of inland farms and retail outlets (11, 13). The data will also provide a solid foundation for future studies on AMR trends in pathogenic *Vibrio* spp. This is important as globally an increase in cases of vibriosis has been observed, and, with global warming, cases are expected to further increase in the future. The AMR data we collected in this study can also be useful in guiding policy development for the non-medical use of antibiotics in farms close to estuaries. In addition, a detailed examination of AMR patterns from clinical strains of *V. parahaemolyticus* isolated during the same period will be useful in exploring if the same declining AMR trends observed for some of the antibiotics, e.g., kanamycin and streptomycin, in food isolates, has also occurred with the clinical isolates.

## MATERIALS AND METHODS

### Bacterial strains

A total of 1021 *Vibrio* isolates, including *V. parahaemolyticus, V. vulnificus, V. fluvialis, V. alginolyticus*, and other *Vibrio* spp. with good identification at the genus level, obtained from wild or cultured molluscs, such as oysters, mussels or clams, were used. The molluscs were harvested from sites on either coast of Canada, and collected between May and October of each year, from 2006 to 2012. All strains were identified, isolated and characterized by standard methods (19, 20). The presence/absence of virulence markers such as thermostable direct haemolysin *(tdh)* and tdh-related haemolysin (trh), were confirmed by polymerase chain reaction [PCR] (19, 21, 22). Standard strains from the American Type Culture Collection (Rockville, MD, USA), i.e, *V. parahaemolyticus* (ATCC 17802), *V. alginolyticus* (ATCC 17749) and *V. vulnificus* (ATCC 27562), were used as positive controls for PCR, and/or phenotypic identification of the strains on selective media. Standard strains of other bacteria *(E. coli* ATCC 25922, *S. aureus* ATCC 25923 and *P. aeruginosa* ATCC 27853) were used as controls in the antimicrobial susceptibility tests, to verify the growth and inhibition zones for each of the antibiotics, as suggested by the ‘Clinical and Laboratory Standards Institute’ [CLSI] (23).

### Microbiological analysis

The antimicrobial susceptibilities of the isolates were determined by the Kirby-Bauer’s disc diffusion method (24) on Mueller-Hinton agar (MHA) plates. In brief, overnight isolated colonies of the test strains from Columbia Blood Agar (Oxoid Ltd., Basingstoke, Hants, UK) supplemented with 5% defibrinated horse blood were re-suspended in tryptic soy broth with 2% NaCl (TSB-2N, pH 8.5), while the (non-vibrio) control strains, grown similarly, were resuspended in trypticase soy broth (TSB; pH 7.4). All suspensions were adjusted to a 0.5 McFarland turbidity standard (ThermoFisher Scientific, Ottawa) and evenly spread onto MHA by using sterile cotton swabs. Overall, 19 antimicrobial discs (Oxoid Ltd, Basingstoke, UK), each containing a known amount of an antibiotic, were laid on a lawn of the bacteria on MHA plates and incubated at 35ΰC for 18-24 h, following the CLSI protocol. Antimicrobial class, antibiotics and the concentrations of the drugs were: AMINOGLYCOSIDES, gentamicin (GEN) 10 μg, kanamycin (KAN) 30 μg, and streptomycin (STR) 10 μg; CEPHEMS, cephalothin (CEF) 30 μg, cefotaxime (CTX) 30 μg, ceftiofur (CTF) 30 μg; FOLATE PATHWAY INHIBITOR, sulfamethoxazole-trimethoprim (24:1) (SXT) 25 μg; LIPOPEPTIDE, polymyxin B (PMB) 300U; MACROLIDE, erythromycin (ERY) 15 μg; PENICILLINS, ampicillin (AMP) 10 μg, piperacillin (PIP) 100 μg; PHENICOL, chloramphenicol (CHL) 30 μg; QUINOLONES, ciprofloxacin (CIP) 5 μg, enrofloxacin (ENO) 5 μg, norfloxacin (NOR) 10 μg, oxolinic acid (OX) 2 μg; SULPHONAMIDE, sulfisoxazole (SF) 300 μg; TETRACYCLINES, tetracycline (TET) 30 μg, oxytetracycline (OT) 30 μg. Isolates were reported as sensitive, resistant, or of intermediate susceptibility to each antibiotic tested based on inhibition-zone size measurement, as recommended by CLSI (23). In the absence of definitive standards of *Vibrio* species for AMR testing, zone diameters of bacterial standard strains, such as *E. coli* ATCC 25922, *S. aureus* ATCC 25923 and *P. aeruginosa* ATCC 27853, were used for quality assurance of the antibiotics, and reliability of the protocol.

### Statistical analysis

Statistical tests (Student’s t-test, and Variance ratio F-test) were used to determine if the AMR datasets from the East and West coasts of Canada were different, or similar. The F-test was chosen to find out the significance of variance in each set from one another; while the t-test analysed the correlation between the east and west coast data related to specific antibiotics. Both tests were done using Microsoft Excel 2010, and a p-value of < 0.05 was defined to indicate significant differences.

## ACKNOWLEDGMENTS

This work was funded (A-base) by Health Canada in support of Canada’s Food Safety Programs. We thank Dr. Franco Pagotto and Dr. Sandeep Tamber of our Bureau (Bureau of Microbial Hazards, Health Canada) for peer-reviewing the manuscript and offering helpful comments. We are grateful to staff from the Media laboratory (Andrew. Samia and Matt) for preparing all the testing materials. We are also thankful for technical assistance from Mahdid Meymandy, Xiaodong Wan and Rachel Adm.

As the study did not involve patient data, the need for informed consent was not required. Also, we declare no competing interests.

